# A low bias method for quantification of Oil Red O staining

**DOI:** 10.1101/2024.06.18.599611

**Authors:** Christian Feldhaus, Veysi Piskobulu

## Abstract

Oil Red O (ORO) is a well established dye for assessing the lipid content of nematodes. Previously reported methods for quantifying ORO staining rely on the difference in absorption in the red part of the spectrum wherORO appears transparent and parts of the spectrum where ORO absorbs light. However, several sources of bias and artifacts are usually not accounted for in previously described methods which makes them unsuitable to detect small differences in lipid content. Here we describe a method which relies on filter based acquisition and semi-automatic image processing including e.g. a correction for different background intensities to eliminate several sources of bias in ORO quantification.

## Introduction

Oil Red O (ORO) is widely used to assess the fat content and distribution in nematodes. However, quantification methods described up to now are prone to bias as they e.g. a) are based on visual scoring (1), b) depend on manual adjustment of analysis parameters (2), c) use image formats which corrupt intensity values (3).

Furthermore, the absorption is often measured by calculating the difference in transmittance between the red (where ORO appears transparent) with the transmittance in a spectral range where ORO absorbs (blue to green range) (4). However, when doing so it needs to be taken into account that by just subtracting the channels there is a potential bias by the different spectral absorbance/ sensitivity of the imaging system due to spectral transmission of optical components or the response curve of the camera (which can be strongly influenced in colour cameras by the set white balance values).

While the mentioned methods are working well for detecting substantial differences in lipid metabolism, smaller differences might get unnoticed or wrongly quantified. Here we present a method for ORO based relative lipid quantification which eliminates several sources of bias by optimising acquisition settings and the analysis pipeline.

## Material and Methods

### Oil-Red-O staining

ORO staining method was modified from (5).

#### preparation of staining solution

Oil-Red-O stock was prepared by dissolving 0.5g ORO (Sigma-Aldrich, product number: O0625) in 100ml isopropanol and subsequent shaking for several days. For preparing ORO working solution the required amont was taken from stock and centrifuged for 5 min at 4500 rpm to get rid of precipitates. Supernatant was diluted with purified water to a 60% working solution which was shaken for 10 min and then centrifuged for 5 min at 4500rpm before usage and only supernatant was applied for staining.

#### Fixation and dehydration of worms

Worms were washed from plates with 1xPBS (phosphate buffered saline), collected in a 15ml tube and supernatant was removed after they sedimented. Worms were transferred to a 1.5ml tube, briefly centrifuged and the supernatat removed. Worms were then fixed in 1% formaldehyed in 1xPBS (freshly prepared from 16% stock) for 30min. Then worms underwent 3 freeze-thaw cycles (each 8min liquid nitrogen followed by tap water thawing) and then 3 times washing in 1xPBS. Samples were then dehydrated in 1ml 60% isopropanol for 2min.

#### staining and mounting

Worms were stained for 30min after applying the staining solution through a 0.22µm sterile filter. Subsequently worms were washed 3 times with 1xPBS and centrifuged. After removal of supernatant they were resuspended in Vectashield (Biozol, catalogue number: H-1000), gently mixed and then mounted on a microscope slide.

### Image acquisition

Images were acquired on an Zeiss AxioImager.Z1 with 100W halogen lamp, EC Plan-Neofluar 10x/0.30 air objective and Axiocam 506 mono camera (Zeiss, Oberkochen, Germany). The 3 channel images were acquired as transmitted light images with channel 1 as normal DIC channel, channel 2 as brightfield channel with Zeiss filterset 49 (DAPI) swapped in the light path and channel 3 as brightfield channel with Zeiss filterset 45 (Texas Red) swapped in the lightpath. Alternatively, an AxioImager.Z1 was used, equipped with Plan-Apochromat 10x/0.45 air objective, and AxioCam 702 mono. Channel settings were identical except a for channel 3, where TxRed ET filterset was used (AHF, Tuebingen, Germany). Light in the transmission range of the DAPI filterset gets absorbed by Oil Red O while the dye is transparent in the transmission range of the Texas Red filtersets.

For all images to be quantitatively compared it is key to keep settings for the lamp intensity, exposure time, objective and filters the same.

During acquisition, care was taken to not have any saturated pixels in the images as those areas would not be quantifiable anymore.

For data saving, storage and analysis only the original data in CZI format was used to make sure all pixel information as well as all metadata are preserved properly.

### Image analysis

Image analysis is automated by running the supplementary script named ORO_measure2.ijm in Fiji (6) (https://fiji.sc/). The script is used as follows:

1. open an image
2. start ORO_measure2
3. when the request pops up, mark a circular ROI (region of interest) for normalising signal (sort of “white balance”).
  a. The ROI
    i. should not touch any worm
    ii. should be away from any image edge
    iii. is ideally somehow next to a worm
4. press “OK” in the pop-up window
5. when the request pops up, mark your ROIs for measurement in the following way:
  a. outline the worm as ROI
    i. be careful not to draw the outline too tight to not exclude signal
    ii. make sure the outline is close enough to the worm to minimise the risk of getting outside false signal (e.g. from dirt) in your measurement
  b. when the outline is closed press “t” to add the worm ROI to the ROI manager of Fiji
  c. if you want to add another ROI go back to a., otherwise proceed
6. press “OK” in the pop-up window
7. if you want to measure another image, close the current image and start again at 1., otherwise proceed
8. close the current image and save the Results table. All measurement ROIs and the modified images are automatically saved in the folder of the original images.

The script performs the following processing steps:

From the area that is first marked outside of worms a ratio for 100% transmittance in both channels gets calculated. By dividing the blue channel by this correction factor differences in the spectral response of the system get compensated as the ratio between the red and blue channel is then normalised. After that normalisation step a new channel is calculated by subtracting the normalised blue channel from the red channel to get the absorbance for ORO.

After outlining the worm area, mean intesity, (raw) integrated density and median intensity for each worm ROI are automatically calculated from the channel containing the absorbance information.

Results are automatically written into a table which also contains information about the image and ROI identity to allow backtracking of the data. The measurement for the absorbance is listed in channel 4.

At the end of the script the processed images and ROI outlines are automatically saved to be able to track each data point in the final results table back to the original raw data. Images are saved as TIFF to preserve full structural and intensity information.

## Results & Discussion

Using a filter based system with a greyscale camera helps to keep the spectral response constant within the system without bias from white balance (which changes the relative sensitvity for the 3 channels present in the Bayer pattern of the camera chip). Bias from white balance can even be present when switching to the greyscale mode of many colour cameras if the white balance settings are not reset before switching. In addition, we use the correction factor to compensate for differences in the spectral response of the system to be able to use the difference or the ratio between the corrected blue channel and the red channel as readout for ORO absorbance. Without this normalisation step you will get a wrong estimate of how much intensity is absorbed by ORO: the intensity in the red and blue channel without any absorbing material in the sample plane is always different (Fig.1), so one will always get an absorbance value even if there is no sample. Without using the correction factor one would need to calibrate the imaging system in a way that the same amount of photons is recorded in an empty sample area in the red and blue channel, which would be hard to achieve and a very tedious procedure. When using a LED lightsource the correction factor will also keep the ratio between the 2 channels constant (Fig.1, this does not work with halogen lamps, data not shown).

**Figure 1:**
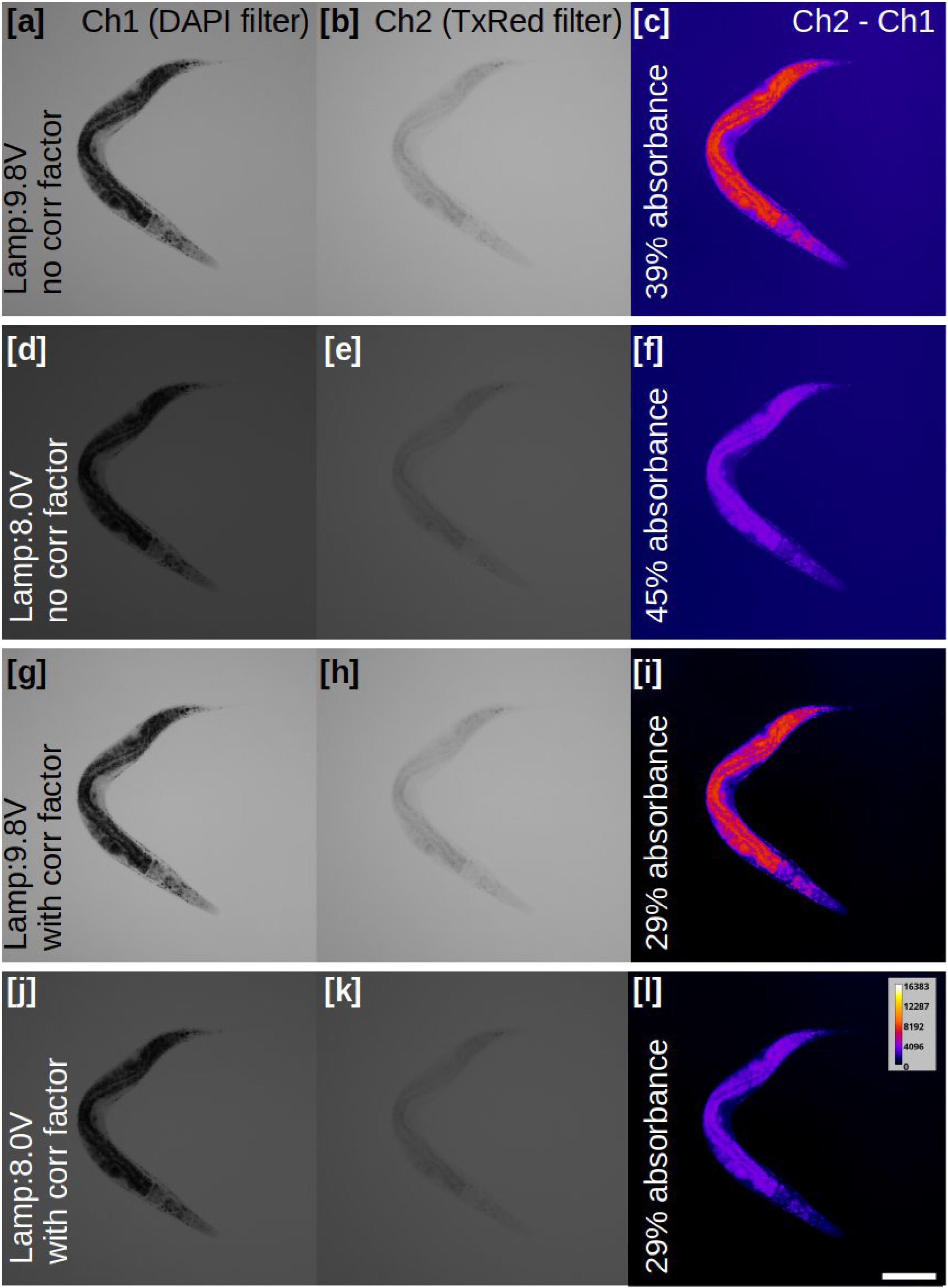
showing the same worm imaged on a system with LED lamp using 2 different lamp intensities and displaying the different absorption values when (not) using the script with the correction factor; note the different background intensities for channel 1 and 2 when not using the correction factor; scale bar is 100µm; colour coding in the right column shows the 14bit range of the acquired images according to calibration bar, left (DAPI filter) and middle (TxRed filter) column are greyscale.

Instead of thresholding the worms we decided for manual outlining as thresholding proved to be unreliable. Frequently the resulting outlines were too tight with signal being excluded. One reason is that standard tresholding algorithms use image statistics as a basis for setting the threshold. This requires to have very similar object distributions and labelling in all your images and it gives very differing results if you have objects of varying number, size or intensity. As the latter is the case for many of the images we usually deal with, thresholding didn’t perform good enough for segmenting worms and we decided for manual outlining.

To assess how much lipids are stained by ORO, the integrated density value for each worm gives the most unbiased readout as it is independent from the marked area. However, with a well-trained researcher marking the outlines or by using carefully adjusted thresholds, the mean might give similarly reliable results with the added value of being able to set the lipid content in relation to worm size.

For the integrated density one can choose to either read out the absolute absorbance in arbitrary units (channel 4 of the processed image) or choose a relative readout for absorbance (channel 4 of the processed image divided by channel 3) or transmittance (channel 2 of the processed image divided by channel 3).

The method presented takes care of several potential sources of bias in ORO quantification. It is therefore suitable for assessing substantial as well as small differences in lipid metabolism using ORO staining.

## Supporting information

Supplementary Script

## Acknowledgements

The authors would like to thank Adrian Streit for bringing up the idea of having a robust method for Oil Red O quantification and Alex Dulovic for help with initial data acquisition. Additional thanks go to the BioOptics Facility at the MPIfor Biology Tübingen and all the people that help keeping the worms happy.

## Author information

## author contributions

CF – conceptualisation, imaging workflow, scripting, main writing

VP – data acquisition, sample preparation, partial writing

Fiji script “ORO_measure2.ijm”:

run(“Set Measurements…”, “area mean integrated median display redirect=None decimal=3”);

nameorig=getTitle();

filepath=getInfo(“image.directory”);

dotIndex = lastIndexOf(nameorig, “.”);

if (dotIndex!=-1) name = substring(nameorig, 0, dotIndex);

rename(name);

run(“Split Channels”);

setTool(“oval”); selectWindow(“C2-”+name);

waitForUser(“Please mark a background ROI.”);

roiManager(“Add”);

getStatistics(areatwo, meantwo, mintwo, maxtwo, stdtwo, histogramtwo);

run(“Select None”);

selectWindow(“C3-”+name); roiManager(“Select”, 0);

getStatistics(areathr, meanthr, minthr, maxthr, stdthr, histogramthr);

run(“Select None”);

corrfactor=meantwo/meanthr;

selectWindow(“C2-”+name); run(“Divide…”, “value=corrfactor”);

imageCalculator(“Subtract create”, “C3-”+name,”C2-”+name);

run(“Merge Channels…”, “c1=C1-”+name+” c2=C2-”+name+” c3=C3-”+name+” c4=[Result of C3-“+name+”] create”);

//run(“Channels Tool…”);

Stack.setDisplayMode(“grayscale”);

roiManager(“deselect”);

roiManager(“delete”);

setTool(“polygon”);

waitForUser(“Please mark your worms. (press [t] to add to ROI Manager)”);

roiManager(“deselect”);

roiManager(“multi-measure measure_all append”);

roiManager(“save”, filepath+name+”_ROISet.zip”);

selectWindow(name);

saveAs(“tiff”, filepath+name+”_M”);

roiManager(“deselect”);

roiManager(“delete”);

